# Genetic Circuits to Detect Nanomaterial Triggered Toxicity through Engineered Heat Shock Response Mechanism

**DOI:** 10.1101/406918

**Authors:** Behide Saltepe, Nedim Haciosmanoğlu, Urartu Özgür Şafak Şeker

## Abstract

Biocompatibility assessment of nanomaterials has been of great interest due to their potential toxicity. However, conventional biocompatibility tests are short of providing a fast toxicity report. We developed a whole cell based biosensor to track biocompatibility of nanomaterials with the aim of providing fast feedback for engineering nanomaterials with lower toxicity levels. We have engineered promoters of four heat shock response proteins. As an initial design a reporter coding gene was cloned to downstream of the promoter regions selected. Initial results indicated that native HSP promoter regions were not very promising to generate signals with low background signals. Introducing riboregulators to native promoters eliminated unwanted background signal almost entirely. Unfortunately, this approach also leads a decrease in expected sensor signal. Thus, a repression based genetic circuit, inspired from HSP mechanism of *Mycobacterium tuberculosis* was constructed. These genetic circuits can report the toxicity of Quantum Dot nanoparticles in one hour with high precision. Our designed nanoparticle toxicity sensors can provide quick reports which can lower the demand for additional experiments with more complex organisms.

## INTRODUCTION

Contrary to its name, heat shock mechanism is a universal process exhibited by cells to any kinds of stress such as heat, osmotic stress, chemicals, ions, or nanomaterials. ^1^ Several transcriptomic analyses indicate that exposure to any stress agents, especially to toxic compounds, prompts changes in gene expression profile, especially genes related to stress response ^2-4^, reactive oxygen species (ROS) metabolic processes ^2^,^3^, DNA damage response ^2^, and cell redox homeostasis ^2^. In order to maintain cellular integrity and survival, nanomaterial exposure (i.e., silver nanoparticles (NPs) ^5^, silica NPs ^6^, quantum dots (QDs) ^7^, or carbon nanotubes ^8^) triggers the production of a set of heat shock proteins (HSPs). ^9^ The HSPs are a sub-group of molecular chaperons; accessory proteins that manage mechanisms crucial for the cell survival and maintenance including protein folding and assembly mechanisms. ^10^ Some chaperones such as Hsp60, Hsp70 and Hsp90 cope with misfolded proteins to refold them properly ^11^, while others, such as clpB (or its eukaryotic homolog Hsp104), Lon and HtrA degrade protein aggregates in cells. ^12,13^ Although HSPR is controlled differently in many organisms, some of the chaperons play a common role in different organisms like Hsp70 which is the major stress related chaperon in bacteria as well as in eukaryotes.

In *Escherichia coli*, main HSP response is conducted by DnaK (Hsp70)-DnaJ (Hsp40)-GrpE machinery. Unlike many protein transcription mechanisms in *E. coli*, HSPs are not under the control of sigma ^70^ factors; a universal subunit of RNA polymerase. Instead, HSPs are controlled by a special stress-inducible subunit, namely sigma ^32^ factor, encoded by *rpoH* gene. ^10,14^ Under steady state conditions, sigma ^32^ level is maintained at constant levels due to its unstable nature; however, after exposure to any stress, sigma ^32^ level is dramatically elevated via improved stability as well as increased synthesis. ^10,15,16^ Sigma ^32^ is regulated by a negative feedback loop controlled by DnaK-DnaJ-GrpE mechanism. ^17^ Accumulation of chaperones in this mechanism holds sigma ^32^ and blocks its activity ^18,19^, leading to degradation of sigma ^32^ by FtsH; a special sigma ^32^ degrading protease. ^20^ Therefore, monitoring of HSP levels in cells can be used as a promising stress indicator.

Nanomaterials are of great interest for their wide range of applicability across many areas from medicine to optoelectronics. Nanoparticles have size-dependent tunable optical and physical properties; which are not usual for bulk materials ^21-23^, nanomaterials are widely used in innovative applications such as in medical diagnostics, drug delivery and targeted photo-thermal therapy. In all of these approaches patients have to be exposed to nanomaterials. ^24^ Also, utilization of nanomaterials in consumer goods may contaminate environment, food and textiles. ^25^ Despite their success in many applications, nanomaterials’ high surface-to-volume ratio indicates potential health problems. In addition, due to their small size, nanomaterials are able to penetrate through cellular barriers easily which may cause cellular stress and many adverse effects such as protein unfolding ^26^, DNA damage ^27,28^, ROS generation ^29-31^, and disruption of gene expression ^27,28,32^. At the system level, nanomaterials can trigger inflammation and alter immune system response ^33-35^. Thus, development of a quick and reliable sensor system that reports nanomaterial-triggered toxicity is very critical for monitoring of biocompatibility analysis of nanomaterials. Specifically, toxicity assessment methods are mostly applicable to chemicals. Indeed, nanomaterials posses some unique properties aside from their chemical composition. Besides the characterization of nanomaterials, their biological uptake mechanism and toxic effect on different pathways should be considered at *in vitro* as well as on *in vivo* models. ^36^ Hence, a quick biosensor system indicating toxicity status of nanomaterials would enable tuning the material properties with further modifications (i.e. surface modifications). Such analysis can give the opportunity to modify nanomaterials immediately by utilizing the feedback from the toxicity tests to increase their biocompatibility.

Toxicity of heavy metals ^37-39^ and other toxic compounds ^40^ to cells have been investigated by many studies ^41^ showing that HSP response is quicker and more straightforward way to evaluate negative effects caused by different stress agents. To observe the HSP response levels of stress agents, construction of a stress sensor circuit with a reporter might help to make comparative analysis over gene expression profiles of cells under some specific stress conditions such as heavy metal exposure. Synthetic biology allows construction of different biosensors with complex genetic circuits to detect analyte of interest. ^42,43^ In this manner, a biosensor circuit coupled with HSP regulatory elements, namely HSP promoters, and a reporter is one of our scopes to detect stress response of cells caused by nanomaterial-triggered toxicity. Many HSP promoters ^4,44,45^ such as dnaK, grpE, clpB, or fxsA have been studied in literature with different reporter genes such as *gfp* ^41,46-48^, *lacZ*, or *lux* ^37,40,46,49,50^ to detect the stress response caused by pollutants and many chemicals.

This study aims to construct a nanomaterial-triggered toxicity sensor (Figure 1a) in order to evaluate biocompatibility of nanomaterials to provide quick feedback about the biocompatibility of the nanomaterials for further engineering and to make the sensor rapid, tightly controlled, easy-to-use, and cheap. For this purpose, we constructed genetic circuits in an engineered cell by employing four promoter sequences (dnaK, clpB, ibpA and fxsA) based on high expression levels of HSPs under stress conditions. First, we tested the heat shock response of toxicity sensors to see how they respond to the stress condition they are evolved for.

**Figure 1.**
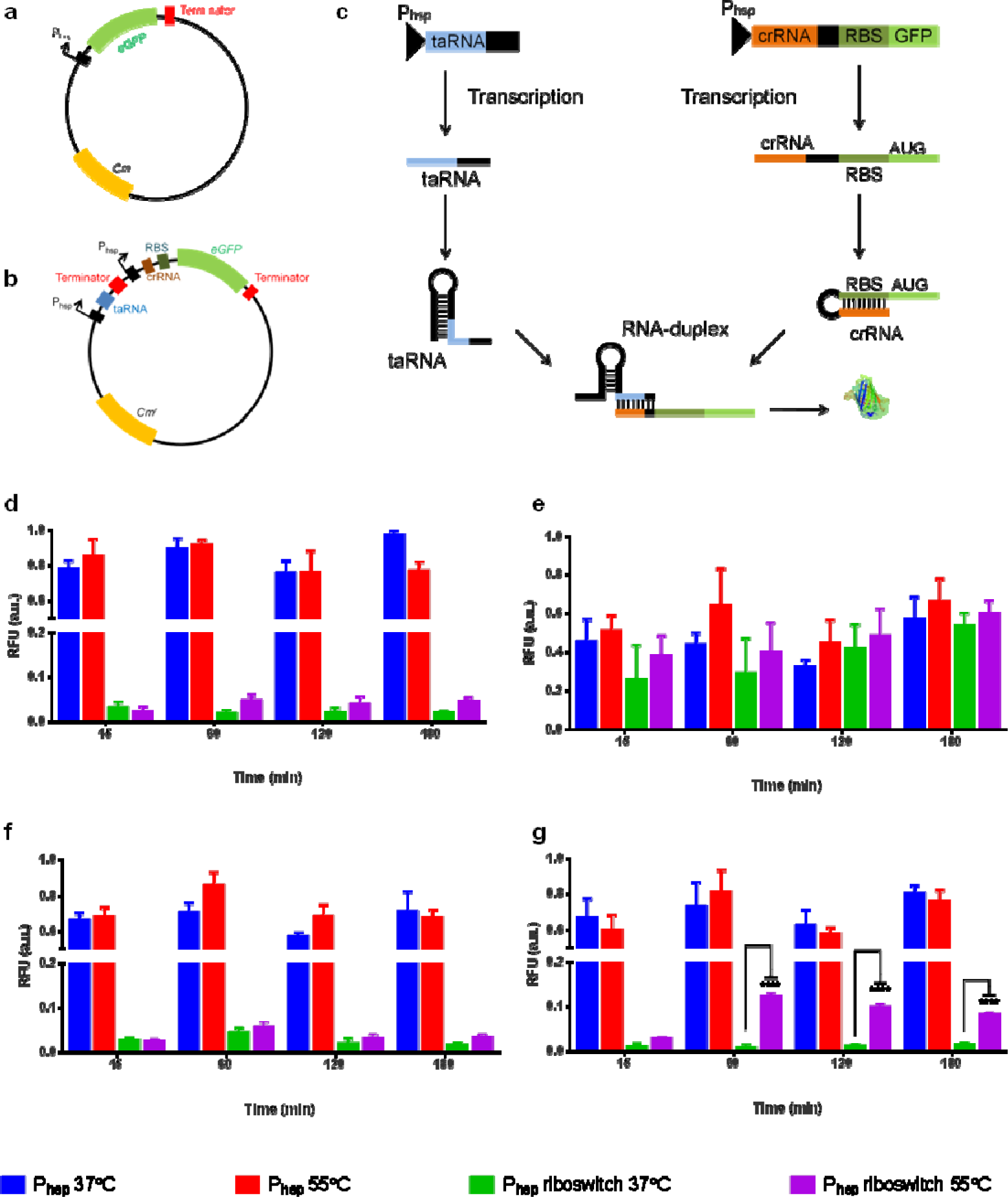
Fluorescence signal results of stress sensors with engineered riboregulators. **a** Representative plasmid map of stress sensors with native HSP promoters. A reporter (gfp) is expressed under the control of heat shock promoters; dnaK, clpB, fxsA or ibpA. **b** Representative plasmid map of stress sensors with engineered riboregulators. Both taRNA and crRNA are controlled by the same HSP promoter. **c** Working mechanism of engineered riboregulators. In the absence of taRNA, reporter expression is blocked by crRNA with a loop formation. However, taRNA favourably forms a complex with crRNA which makes RBS free so that gene expression starts. Fluorescent signal results of heat treated toxicity sensors with native HSP promoters and their riboregulator-mediated constructs for P _dnaK_ **(d)**, P _clpB_ **(e)**, P _fxsA_ **(f)**, and P _ibpA_ **(g)** circuits. Experiments were performed as three biological replicates in different days. Heat shock was applied at 55°C water bath for 30 min, and control samples were kept at 37°C. Sensors with native HSP promoters and sensors with riboregulators in each group were normalized between each other based on formula stated in Materials and Methods section. *p* ≤ 0.0001 is represented with four stars while statistically non-significant results have no stars.

Secondly, we modified our toxicity sensor circuits with riboregulators ^51^ to eliminate any leakage may be caused by HSP promoters. Isaacs *et al.* and characterized a library of artificial riboregulators to tightly control the gene expression. This system employs small non-coding RNAs (sncRNAs) found in cells. Based on this strategy, cis-repressing RNA (crRNA) blocks gene expression via forming loop on ribosome-binding site (RBS). However, upon induction, trans-activating RNA (taRNA) favourably forms complex with crRNA releasing RBS free so that ribosome can recognize RBS sequence and initiate gene expression (Figure 1c). The riboregulators constructed by Isaacs *et al.* were inserted to HSP promoters at transcription initiation sites of toxicity sensors (Figure 1b). Riboregulator-mediated toxicity sensors showed a dramatic decrease in background signal.

In addition to the HSP promoters we exploited from *E. coli* for the circuits, we employed a transcription repressor found in HSP pathway of *Mycobacterium tuberculosis* to build a hybrid sensor system. ^52^ We engineered natural *E. coli* dnaK promoter with repressor binding regions (IR2 and IR3 sequences) from natural *M. tuberculosis* dnaK promoter to construct toxicity sensor circuits. Results showed that IR2 and IR3 sequences help repression of the signal almost entirely under steady state conditions. Formed circuits were tested by exposing them to the QDs for a toxicity assay.

In this work we have demonstrated the utilization of engineered heat shock response mechanism as a powerful candidate to build ordered genetic circuits to report nanomaterial triggered toxicity. Such a biological device can serve as a quick analytical setting to provide initial data about the cytotoxicity of the nanomaterials of interest without any need for a more complex assay. This initial data can be used as a feedback to engineer or modify the nanoparticles.

## RESULTS AND DISCUSSION

### Characterization of Stress Sensors via Heat Shock

Selection of a proper promoter is crucial in whole cell biosensor studies since reporter expression rate highly dependents on promoter strength. Heat shock response pathway offers many promoter options with varying strengths from very low to high expression level. Thus, we chose HSP promoters which show moderate or high expression levels in response to heat induction. ^4^ We considered following heat shock promoters which were characterized in previous studies namely: dnaK ^4,41,44,45^, clpB ^41,44,45^, fxsA ^4^ and ibpA ^4^.

We constructed our stress sensor plasmids with *gfp*, which is expressed under the control of HSP promoters; dnaK, clpB, fxsA, and ibpA, the plasmid maps of these initial designs can be found in Figure 1a, 1b. As a consequence of the leaky nature of the primary circuits formed, we engineered primary circuits with riboregulators adapted from Isaacs *et al.* ^*^51^*^ The initial test results of the primary circuits with increased heat stress are presented in in Figure 1d, 1e, 1f, 1g. Riboregulator-modified and non-modified primary sensor circuits are compared.

The native HSP promoters are active in cells while the cells are growing. They trigger a notable signal even at 15 min corresponding to time zero. To be able to make a significant comparison, 15 ^th^ minute was used as the starting point. This has also allowed us to eliminate any errors due to the delay in sampling and measurements. In general, reporter expression reaches its maxima at 1 hour after heat induction and the signal decreases afterwards which might show that cells start adapting themselves to the environment and decreasing sigma ^32^ - dependent gene expression. Besides transcription, decrease in translation efficiency might cause decline in signal accumulation. ^4^ Thus, not only promoters but also RBS strength might be engineered to overcome insufficient signal output.

Among the four primary circuits, the one exploits the clpB promoter region (originally active in the synthesis of the ClpB heat shock protein) did not give a useful signal. However, rest of the primary circuits gave a high signal both in the presence and absence of the stress condition, which was the elevated temperature. But the increase in the signal upon heat exposure was not significant and a fold change was not notable. That is why; we employed riboregulator systems to prevent any background signal in primary circuits. Riboregulator-mediated circuits have a dramatic effect on turning down the background signal, except for clpB promoter. ClpB promoter did not show any significant effect on removing background signal.

Among the other promoters, the level of the background signal was very similar after the addition of the riboregulators. Nevertheless, riboregulator modified ibpA promoter showed the best performance compared to the DnaK promoter, ClpB promoter and fxsA promoter upon exposure to the elevated temperature stress.

The confocal images on Figure 2 showed the responses of the genetic circuits upon stress exposure, visualizing the effectiveness of the riboregulators. However, one should also point that the riboregulators also turn down the background signal while they provide an obvious difference between toxicity exposed and unexposed cellular devices.

**Figure 2.**
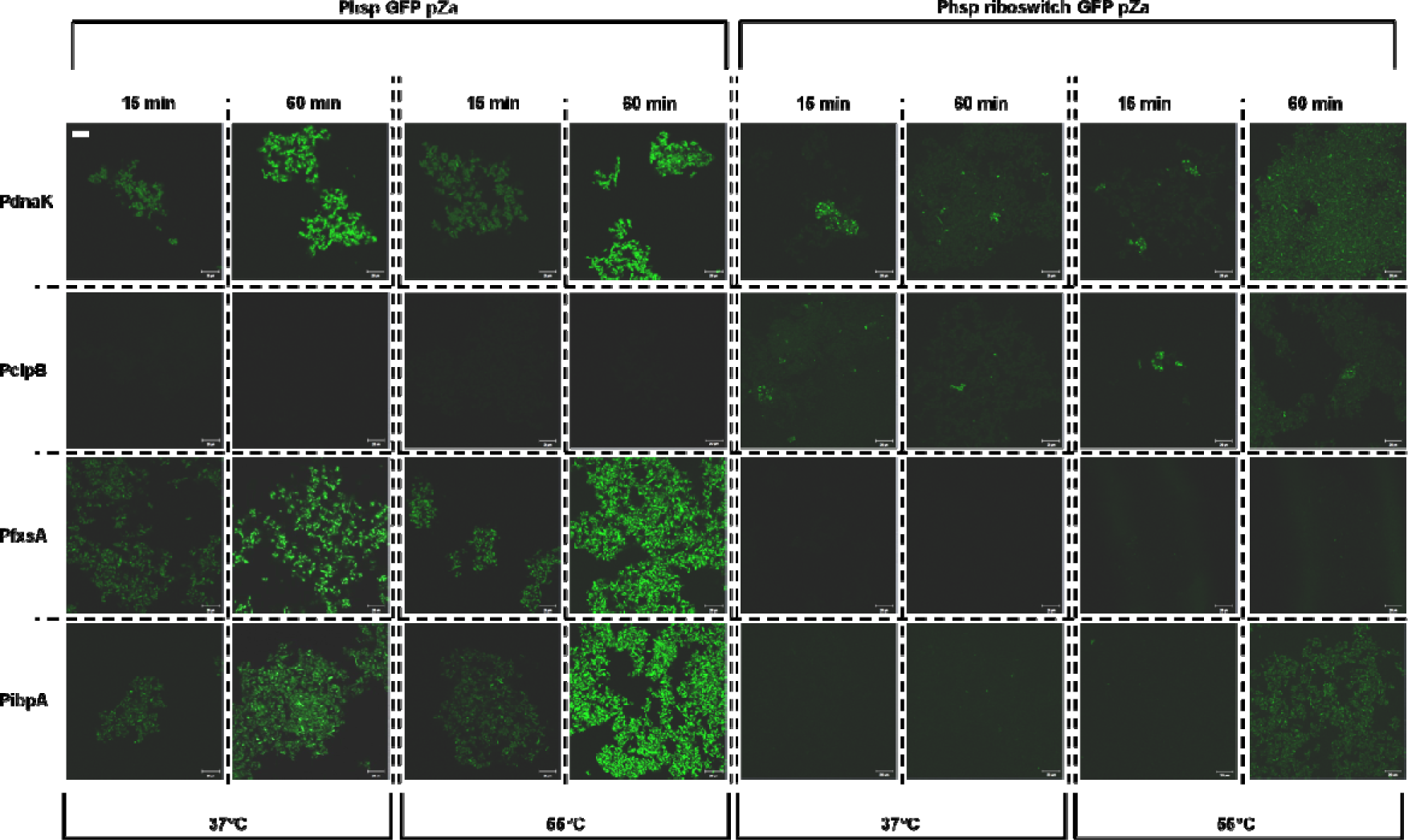
Time dependent confocal microscopy images of heat shock treated primary toxicity sensors. Each row represents stress sensors with HSP promoter (left) and its riboregulator-mediated construct (right) (P _dnaK_, P _clpB_, P _fxsA_ and P _ibpA_, respectively). Each column indicates fluorescence of stress sensors upon heat treatment at 37°C and 55°C at 15 min and 60 min which are the first-time point and the highest signal point of stress sensors, respectively. Scale bar located top left corner indicates 20 µm and valid for each representative image found in this figure.

### A Repressor-based HSP Control of Stress Sensors

HSP mechanism shows differences among prokaryotes. Unlike HSP mechanism in *E. coli, M. tuberculosis* regulates its HSP mechanism with a special repressor, so called HspR, synthesized from dnaKJE-hspR operon via a self-controlled feedback mechanism. HspR recognizes special sequences found in promoter region called HspR-associated inverted repeats (HAIR) with the assistance of DnaK chaperon and blocks its own operon under steady state conditions. However, upon stress, HspR and DnaK dissociates from promoter initiating the gene expression. ^53,54^ Native HSP system of *E. coli* has transcription factors those are working as an activator in the system, which might be the reason why we observed a high background signal in above explained circuits. We proposed that a repression-based sensor design might be a solution to suppress high background signal. This hypothesis led us to design circuits with HSPR systems from *M. tuberculosis*.

We used inverted repeat motifs (IR2 and IR3) found in HAIR sequences ^55^ on *M. tuberculosis* dnaK promoter (mtP _dnaK_) to replace native *E. coli* dnaK promoter (P _dnaK_). Plasmids were constructed with single and double repeats of IR motifs locating at downstream of P _dnaK_. HspR was cloned under a constitutive promoter, proD, and cloned onto another plasmid. Proposed mechanism is to block reporter expression via constitutively expressed HspR under steady state conditions (Figure 3a) while HspR dissociates from the promoter driving reporter expression upon stress (Figure 3b). A comparison on the repression strengths of HspR on native mtP _dnaK_ containing circuits and native P _dnaK_ promoter containing circuits have been shown in Figure 3. Additionally, inducible HspR expression in *E. coli* was demonstrated in Western Blot analysis (Figure S24).

**Figure 3.**
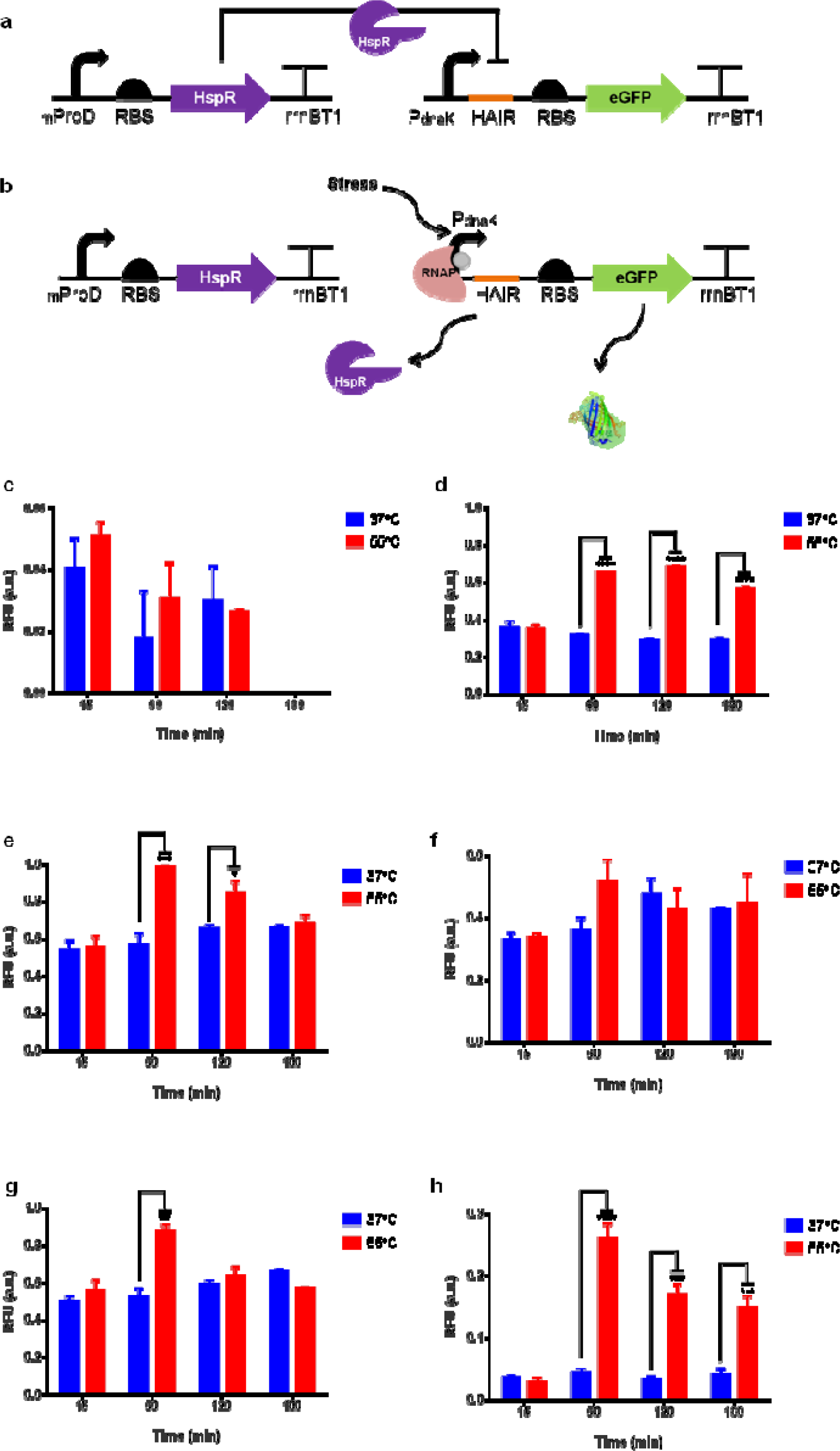
Response of the genetic circuits upon heat shock exposure dnaK promoter with HAIR sequences co-expressed with HspR repressor. Working principle of HspR is shown in **a** and **b**. At steady state conditions, constitutively expressed HspR recognizes specific sequences on promoter region blocking the gene expression **(a)** while releases those specific sequences upon stress exposure initiating gene expression **(b)**. Fluorescence signal results of repression-based stress sensors with mtP _dnaK_ **(c)**, P _dnaK_ **(d)**, P _dnaK-IR2_ **(e)**, P _dnaK-IR3_ **(f)**, P _dnaK-IR2-_ IR2 **(g)**, and P _dnaK-IR3-IR3_ **(h)**. Experiments were performed as three biological replicates in different days. Heat shock was applied at 55°C water bath for 30 min, and control samples were kept at 37°C. Fluorescence intensity of each group was compared with each other and normalized according to formula stated in Materials and Methods section. p ≤ 0.05, p ≤ 0.01, p ≤ 0.001, and p ≤ 0.0001 is represented with one, two, three, and four stars, respectively. Statistically non-significant results have no stars.

Results confirmed that HspR favourably bound to mtP_dnaK_ promoter region and repressed the gene expression. Yet, unlike our expectation, heat treatment was not enough to turn on the gene expression completely (Figure 3c). Besides, HspR repressed native *E. coli* P_dnaK_ less favourably compared to its native promoter, mtP_dnaK_. It is proposed that HspR dissociated from promoter upon heat exposure which led a two-fold increase after 60 minutes (Figure 3d). Compared to the engineered P_dnaK_ sensors with HAIR motifs, single IR motifs showed no or little significant difference in terms of gene expression upon stress exposure (see Figure 3e for IR2 and Figure 3f for IR3). These results confirmed that HspR favourably repressed the gene expression with the help of IR2 and IR3 sequences in the circuits. Meantime, these sequences might trigger a lower degree of dissociation of HspR from promoter so that gene expression could not begin through stress promoters even though there was a stress as an input. Also, double IR2 repeat indicated the same gene expression pattern with single IR repeats upon heat exposure (Figure 3g). On the other hand, double IR3 repeat significantly enhanced HspR repression of P_dnaK_ blocking the gene expression under steady state conditions. It also altered gene expression upon heat showing the highest reporter expression-fold compared with other modifications (Figure 3h). Provided data is supported by microscopy results in Figure 4.

**Figure 4.**
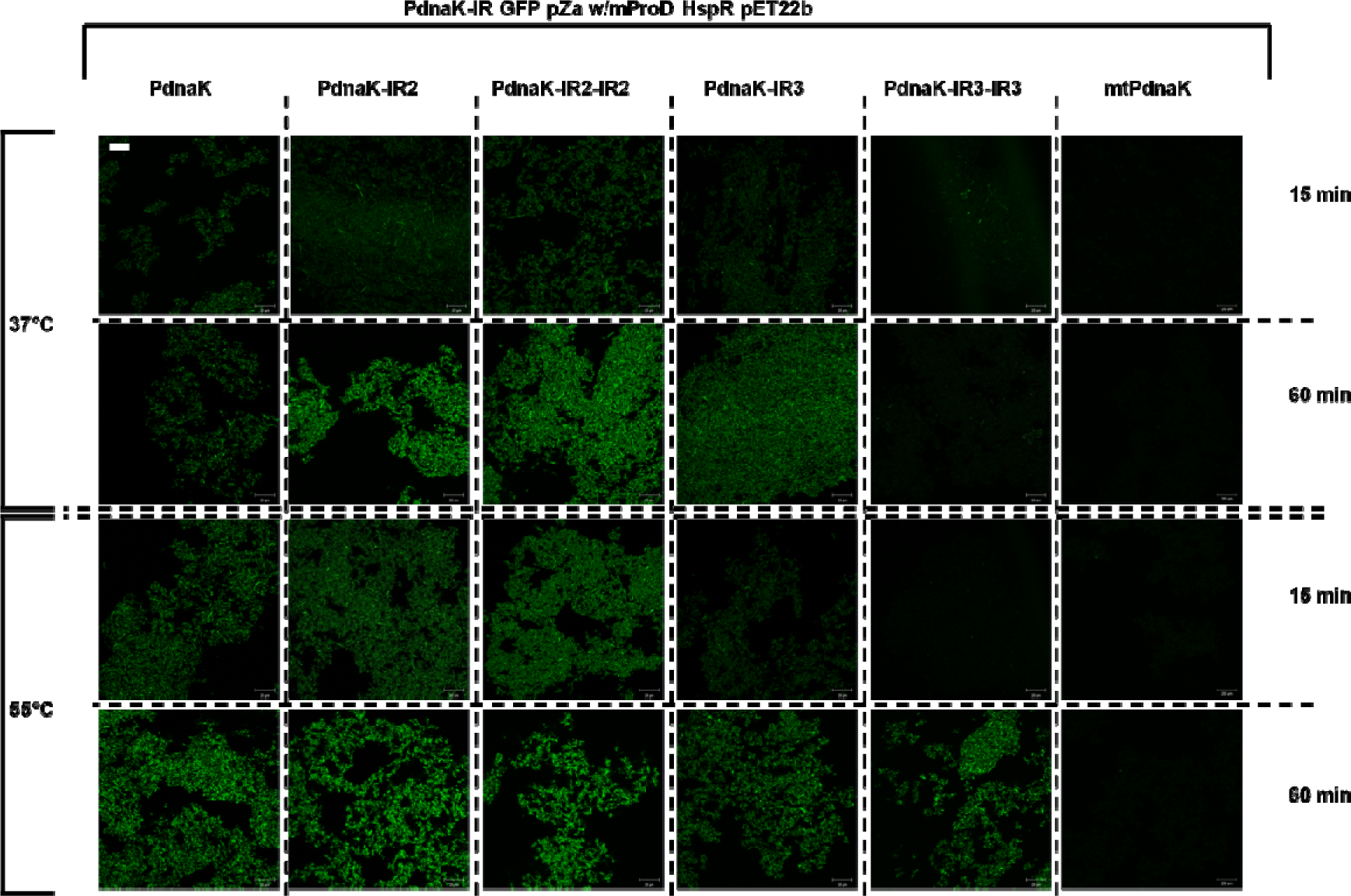
Time dependent confocal microscopy images of modified dnaK promoter with HAIR sequences co-expressed with HspR repressor. Each column represents different modifications of promoters (P_dnaK_, P_dnaK-IR2_, P_dnaK-IR2-IR2_, P_dnaK-IR3_, P_dnaK-IR3-IR3_, and mtP_dnaK_, respectively.). Each row indicates fluorescence of stress sensors upon heat treatment either at 37°C (upper) or 55°C (lower) at 15 min and 60 min which are the first time point and the highest signal point of stress sensors, respectively. Scale bar located top left corner indicates 20 µm and valid for each representative image found in this figure.

As a conclusion, double IR3 sequence plays a strong role in HspR recruitment and turning the promoters ON/OFF. Data supports that the HAIR motifs are good candidates to be used in the designs of the genetic circuits to monitor the stress exposure. HspR toxicity sensors have a lower background signal under steady state conditions while gene expression might dramatically increase upon stress exposure.

### Sensing the Nanomaterial Triggered Toxicity

Among nanomaterials, QDs are used for many applications such as fluorescent labelling or drug delivery since they indicate high photostability also are easy to functionalize. ^56^ Due to the high demand for the utilization of QDs we aimed to use them as potential test materials for their toxicity. Testing nanomaterials for their cytotoxicity is a common approach in every synthesis work for biomedicine related applications. ^57^ A fast feedback about the nanomaterial triggered toxicity can be obtained from the whole sensor system we are proposing here. Such information can help one to engineer nanoparticles and save time without carrying out complex tests at every step. Additionally, such a system can be successfully used to monitor nanoparticle triggered toxicity on environments as well. QDs show toxicity on bacteria with couple of proposed possible ways including photogeneration and ROS formation. Direct release of heavy metals on QD surface caused by light might increase heavy metal ion uptake by cells which cause DNA damage, loosen membrane integrity, interrupted electron transfer chain or oxidation of proteins and lipids in cells. ^30,56^ To analyze nanomaterial-triggered toxicity, two sets of selected stress sensors were screened via red emitting CdTe QDs. The first set, riboregulator-mediated sensors, showed that stress response to higher concentrations (300 nM) appeared quickly at which is right after QD exposure (Figure 5) while lower concentrations (50 nM) was not enough to trigger stress response dramatically. Results were supported by confocal microscopy in Figure 6. Since QD toxicity is concentration dependent, higher QSs in the media might cause higher cellular uptake or ROS generation which trigger stress response much quicker and higher than lower concentrations. Thus, nanotoxicity sensor signal is higher in each case independent from the riboregulator-mediated HSP promoter type. On the other hand, signal start decreasing upon cell death as a result of higher nanoparticle concentration.

**Figure 5.**
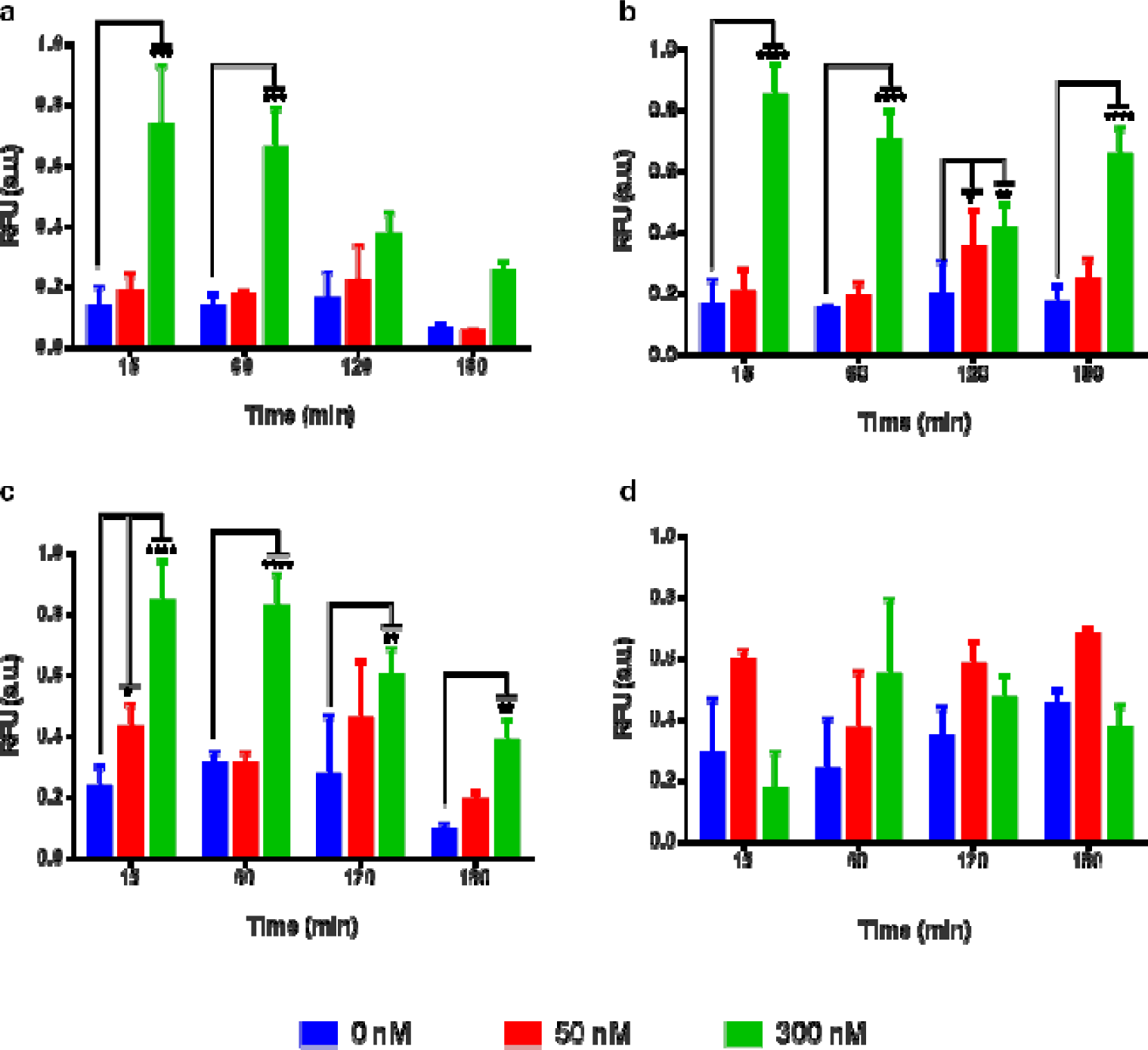
Fluorescense signal results of CdTe QD treated riboregulator-mediated stress sensors with P_dnaK_ **(a)**, P _clpB_ **(b)**, P _fxsA_ **(c)**, and P _ibpA_ **(d)**. Experiments were performed as three biological replicates in different days. QD treatment was applied 50 and 300 nM. All data were normalized according to formula stated in Materials and Methods section. p ≤ 0.05, p ≤ 0.01, p ≤ 0.001, and p ≤ 0.0001 is represented with one, two, three, and four stars, respectively. Statistically non-significant results have no stars.

**Figure 6.**
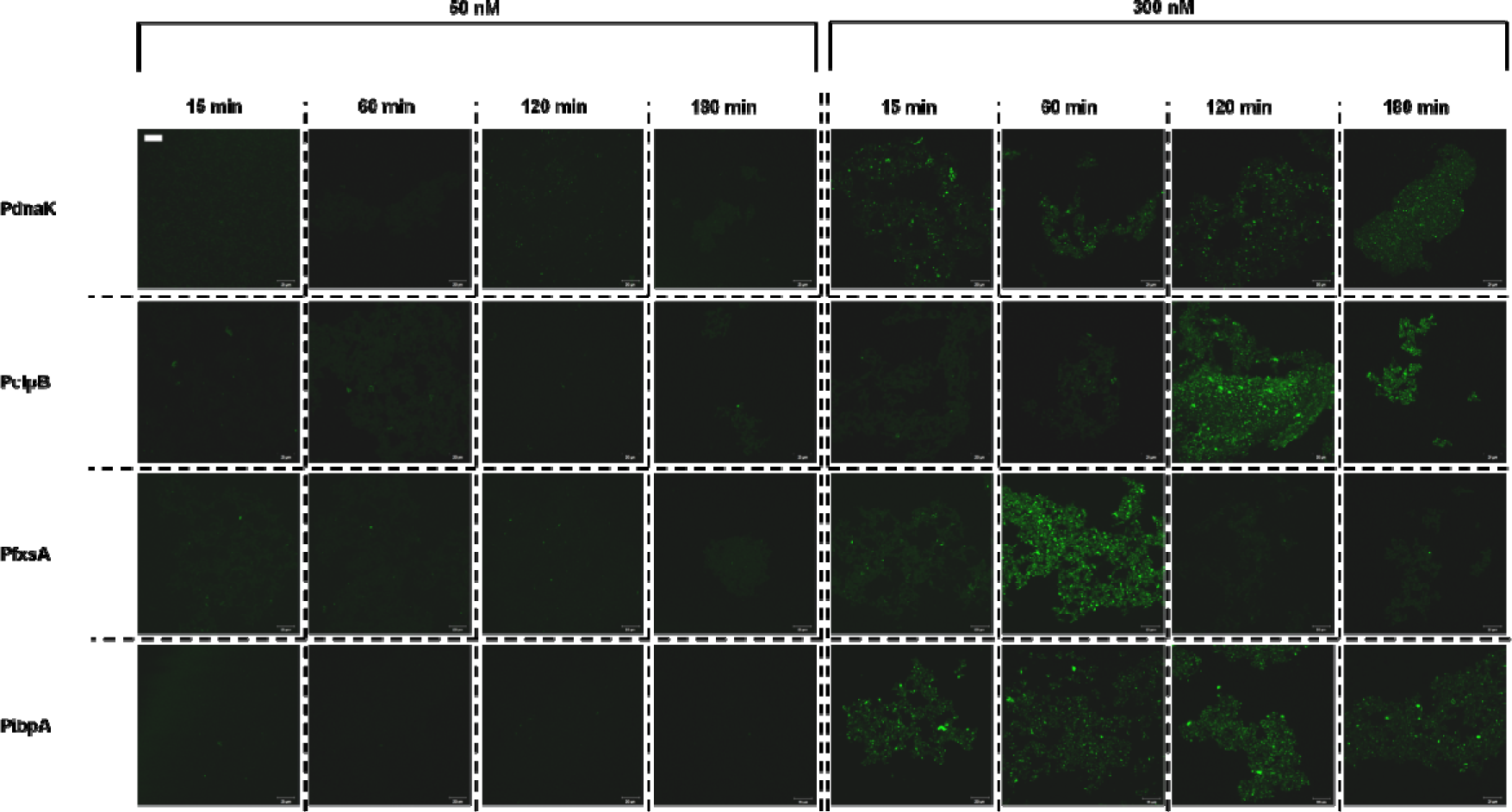
Time dependent confocal microscopy images of QD treated riboregulator-mediated sensors. Each row represents different promoters (P_dnaK_, P _clpB_, P _fxsA_ and P _ibpA_, respectively) and each column indicates time dependent fluorescent response caused by 50 nM (left) and 300 nM (right) CdTe QDs. Scale bar located top left corner indicates 20 µm and valid for each representative image found in this figure.

HspR-mediated sensors generated similar results with the first set (Figure 7). They also showed quick and high response to higher QD concentration. On the other hand, sensor with mtP_dnaK_ shows no response indicating strong repression by HspR that does not allow transcription initiation of the reporter. Besides, overall signal upon induction was higher than the riboregulator-mediated sensors making them good candidates to determine nano-toxicity. The images of the engineered cells containing synthetic genetic circuits were recorded and presented in Figure 8.

**Figure 7.**
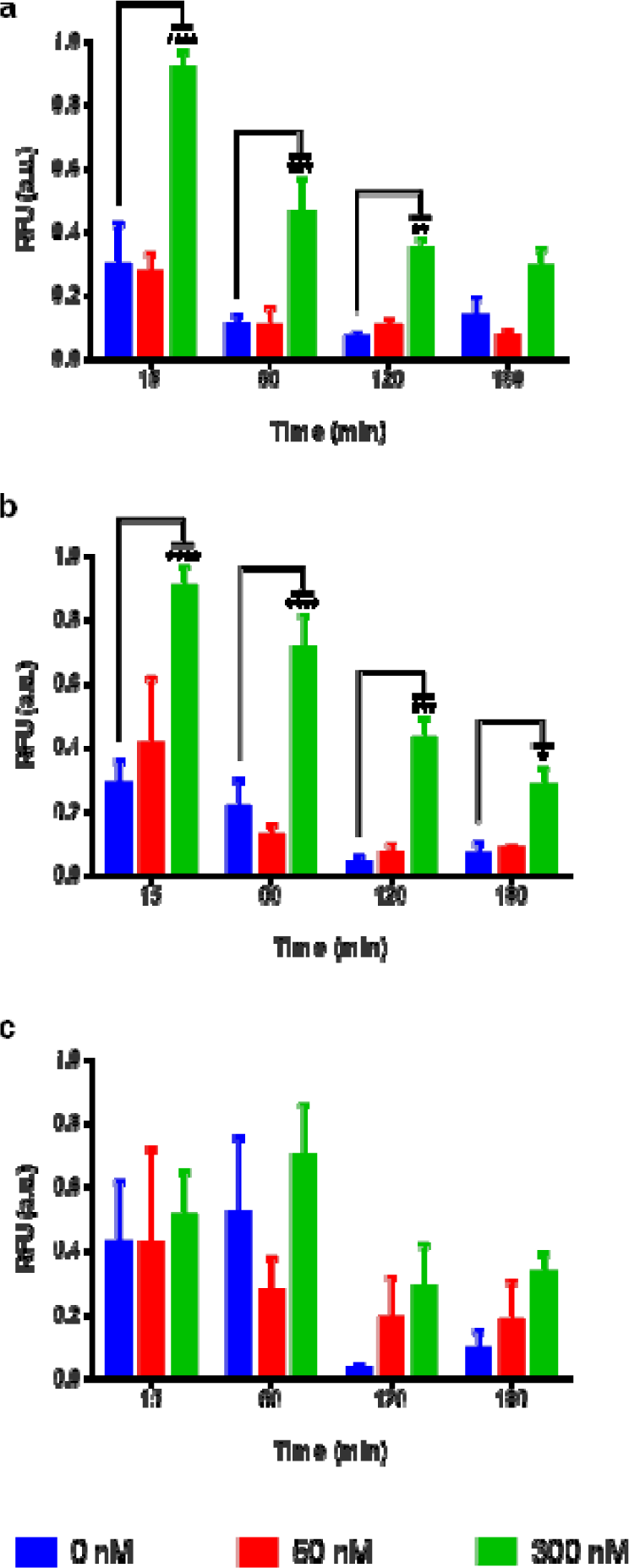
Fluorescence signal results of CdTe QD treated HspR-mediated stress sensors with P_dnaK-IR2-IR2_ **(a)**, P_dnaK-IR3-IR3_ **(b)**, and mtP_dnaK_ **(c)**. Experiments were performed in three biological replicates on different days. QD treatment was applied 50 and 300 nM. All data were normalized according to formula stated in Materials and Methods section. p ≤ 0.05, p ≤ 0.01, p ≤ 0.001, and p ≤ 0.0001 is represented with one, two, three, and four stars, respectively. Statistically non-significant results have no stars.

**Figure 8.**
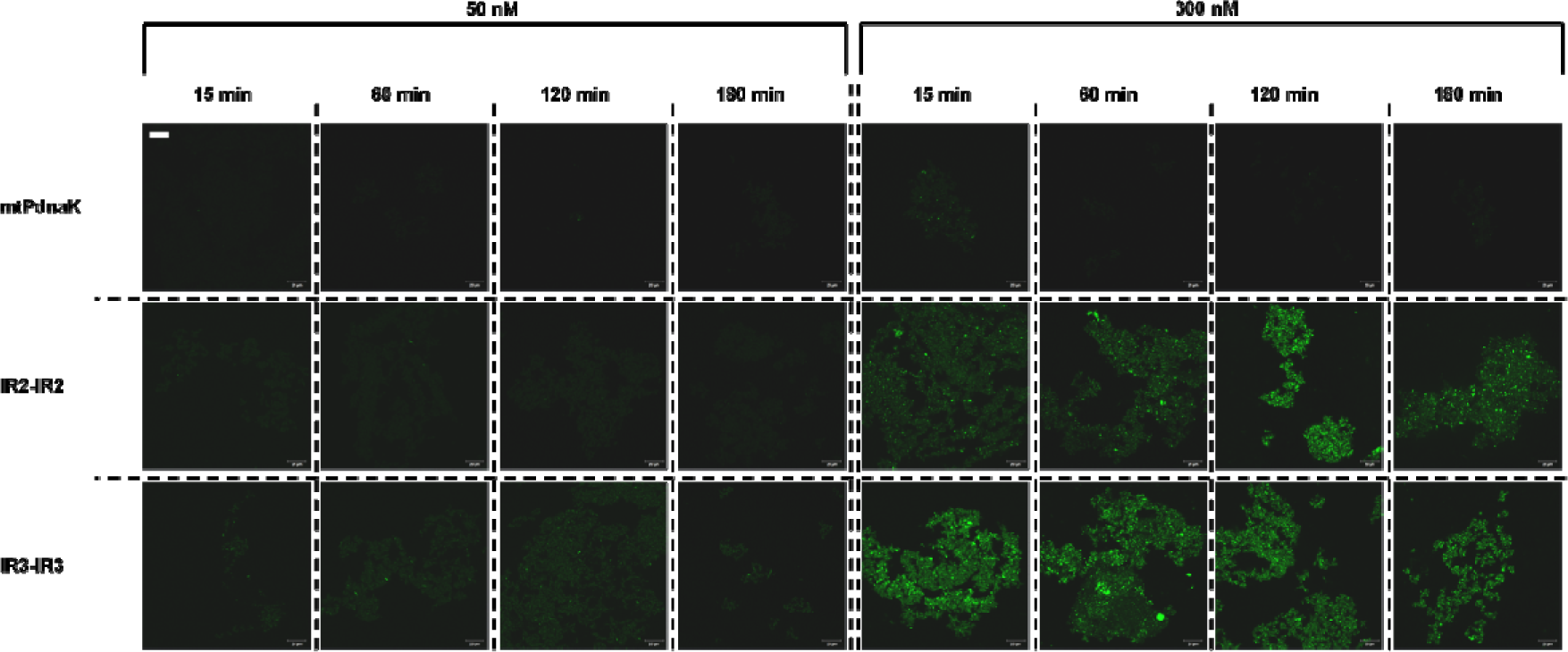
Time dependent confocal microscopy images of QD treated HspR-mediated sensors. Each row represents different promoters (mtP_dnaK_, P_dnaK-IR2-IR2_, and P_dnaK-IR3-IR3_, respectively.). Each column indicates time dependent fluorescent response caused by 50 nM (left) and 300 nM (right) CdTe QDs. Scale bar located top left corner indicates 20 µm and valid for each representative image found in this figure.

To carry out an experiment regarding the gene expression at transcription level one of the best performing set of native HSP promoter coupled with riboregulator, ibpA stress sensor was selected. Using real time quantitative-PCR (RT-qPCR) a representative set of experiment was carried out (Figure S23a). Results showed that gene expression was higher upon heat treatment (55°C) while QD treatment had no dramatic effect compared with untreated control group. Likewise, P_dnaK-IR3-IR3_ sensor was tested to represent HspR-mediated stress sensor group in order to analyse gene expression at transcription level (Figure S23b). A change in gene expression level in agreement with the previous observations was noted.

Following QD characterization with selected sensors, we chose HspR-mediated P_dnaK-IR3-IR3_ sensor as potentially the best nano-toxicity determinant. Further, we analyzed dynamic range of this sensor via CdTe QD treatment (Figure 9). Induction concentrations were determined at log-scale from 0-to-1000 nM of QDs. Results indicated that lower concentrations were not enough to trigger stress, while very high concentrations might cause cell death or signal quenching by internalized QDs. However, stress caused by medium-range concentrations could be observed clearly (Images of these cells can be found in Figure S22).

**Figure 9.**
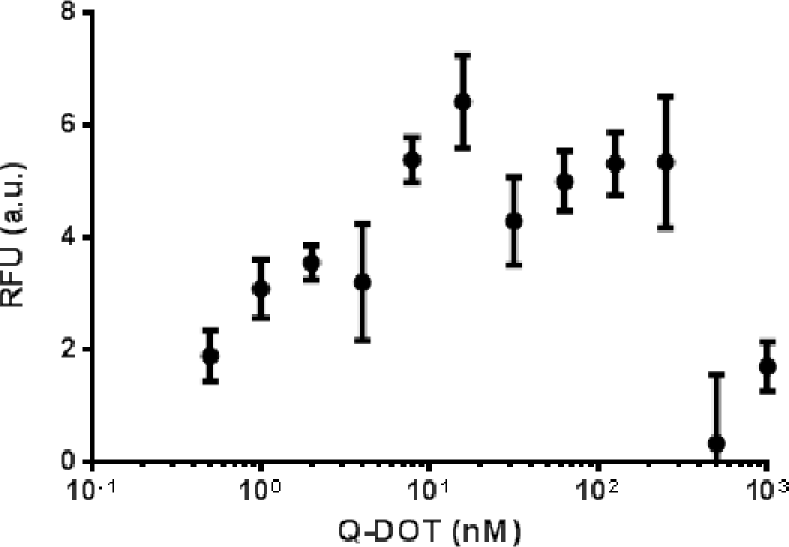
Dose-response curve of selected HspR-mediated sensor (P_dnaK-IR3-IR3_) against CdTe QD treatment. QD concentration range was selected based on log-scale. Experiments were performed as three biological replicates in different days. Measurement was taken at 60 min. after QD treatment.

Toxicity response to nanomaterials is highly cell and chemical composition type dependent. Membrane structure and composition affects nanomaterial uptake through cells. For instance, even Gram negative and Gram positive bacteria do not respond similarly to the same nanomaterial since they have different membrane composition. Besides, lack of LPS makes Gram negative cells more vulnerable against nanotoxicity. ^56^. Here we demonstrate that our engineered bacterial nanomaterial-triggered stress sensors are able to sense and respond accordingly to the stress conditions. Considering that HSPR mechanism is common in each cell type, we expect to transfer and modify this bacterial system to other cell types. Still, cell type response might require some further optimization to tune gene expression based on nanomaterial concentration and type. However, in general, we proposed a whole cell sensor that can produce quick information about the toxicity of the nanomaterials of interest from a global perspective. On the other hand, the proposed sensor does not provide detailed information about the nanomaterial-triggered toxicity including its action mechanism at the downstream of pathways of interests.

## CONCLUSION AND FUTURE PERSPECTIVES

Recent developments in nanotechnology accelerated nanomaterial applications in various fields. Their unique properties ^21-23^ such as small size, high surface-to-volume ratio, catalytic activity make them potentially dangerous to living systems because they have ability to penetrate tissues and cells easily. ^26-35^ Thus, an early diagnostic toxicity assessment procedure is a necessity prior to nanomaterial application on living systems as well as on environment. Here we proposed synthetic genetic circuits which are capable of sensing early stages of stress caused by nanomaterials using an engineered HSPR system. ^9^ In this study, all stress responsive circuits were characterized by exposing the circuit harbouring cells to elevated temperatures, later selected cells were used as the candidate sensors for nanomaterial exposure. Stress responsive circuits with mostly expressed native HSP promoters controlling a reporter output, GFP. Native HSP promoters were found to have a looser control on the expression and as a result high background signals were observed. Coupled with a set of synthetic riboregulators, engineered native HSP promoters were found to be more functional as sensing elements. Among the native promoters chosen, P _ibpA_ promoter also seemed to be the best performing promoter when coupled with a riboswitch system. One should consider that the riboswitch we used prevented leaky protein expression as expected. In addition to the native HSP system in *E. coli*, we searched for other HSP systems in other bacteria. Interestingly, *M. tuberculosis* has a repression based where *E. coli* has an activation based HSPR system. Thus, we employed *M. tuberculosis* system and designed a circuit to sense the nanomaterial triggered toxicity. In this approach, we did not need to employ a riboswitch system as the circuit itself has a repressor-based operation preventing high degrees of protein expression leakage.

Native HSP system is leaky in terms of synthesizing the HSP proteins which has biological basis as the cells are continuously bearing stress conditions during growth. A basal level of DnaK protein, for example, is crucial for the cell to keep its proteins functioning. However, for synthetic biology applications, the functionality of the cells and circuits should be programmed so that the native/non-native systems can be adapted for certain needs. Whole cell sensors have a great potential for numerous future applications monitoring nanomaterial-triggered toxicity is among those. In general, nanomaterial-triggered toxicity is a complex phenomenon. After the nanoparticles enter the blood stream, or interact with the cells many molecular mechanisms are triggered. However, to track these changes at genome and proteome level is laboured intensive and costly. That is why we believe that our quick reporter systems will provide crucial initial data to make judgements about the level of toxicity. However, one should notice that our proposed circuit design is not tissue or organism specific but gives a general idea if the nanomaterial of interest triggers any toxicity. To be more specific about the reasons for the toxicity, specific biomarkers should be identified using genome, transcriptome, or proteome level analysis for each type of nanoparticle available with varying surface properties. Such an attempt may have a potential to develop whole cell sensor with complex circuit designs including logic-based operations.

## MATERIALS AND METHODS

### Media and Strains

*E. coli* DH5α (NEB) was grown in LB medium (1% (w/v) tryptone, 0.5% (w/v) yeast extract, 0.5% (w/v) NaCl) with proper antibiotics at 37°C and 180 rpm shake in Erlenmeyer flasks. Overnight cultures were prepared from frozen glycerol stocks and incubated for 16 h with the same culturing conditions mentioned previously. 1% of inoculums from overnight cultures were used to start experimental culture and monitored via spectrophotometer (GENESYS 10 Bio, Thermo Scientific) until OD _600_ reaches 0.4-to-0.6 before induction steps were applied.

### Plasmid Construction

*E. coli* heat shock promoters were amplified from *E. coli* DH5α genomic DNA while *M. tuberculosis* dnaK promoter was amplified from *Mycobacterium bovis* genomic DNA, which has the same promoter sequence with *M. tuberculosis*, by primers Table S1. Codon optimized *M. tuberculosis HspR* repressor and engineered riboregulators with PclpB promoter part and PibpA-taRNA part were synthesized by GENEWIZ Company. Q5 Hot Start High-Fidelity DNA Polymerase (New England Biolabs, Inc.) was used for all PCR reactions. To construct the stress sensor plasmid backbone, pZa-tetO-eGFP vector was digested with XhoI-KpnI restriction enzymes (New England Biolabs, Inc.) and pET-22b(+) was digested with SalI-SpeI restriction enzymes (New England Biolabs, Inc.) for repressor plasmid backbone construction (pET22b-mProD-HspR). NucleoSpin Gel and PCR Clean-up kit (Macherey-Nagel) was used according to manufacturer’s instructors to purify digested DNA samples or PCR products from 1-1.8% Agarose gels stained with SYBR Safe DNA Gel Stain (Thermo Fisher Scientific). Plasmid construction was made via ligation with DNA T4 ligase (New England Biolabs, Inc.) or via Gibson Assembly method describe by Gibson et al. ^58^. Expression vector for HspR (T7-HspR-pET22b) was constructed via Gibson Assembly method. First backbone was digested with XbaI-XhoI enzymes and HspR was amplified at two round PCR with specified primers in Table S1. After all assembly methods, mixes were directly transformed into chemical competent *E. coli* DH5α cells. Colonies were screened by plasmid digestion. Constructed genetic circuits were sequence verified by Sanger Sequencing (GENEWIZ).

### Heat Shock Experiments and Toxicity Assay

After OD _600_ of experimental cultures reached 0.4-to-0.6, Erlenmeyer flasks corresponding to heat shock treatment were immersed in 55°C water bath for 30 minutes as describe by Rodrigues et. al ^4^ while Erlenmeyer flasks corresponding to toxicity assay were treated with water-soluble thiol-capped CdTe QDs with varying concentrations. QDs were prepared using the method as explained in a previous work Seker et al. ^59^ Control flasks were kept as untreated at 37°C. Three independent biological replicas were prepared for each group.

### Fluorescence Measurement and Data Analysis

All fluorescence measurement studies were conducted via microplate reader (SpectraMax M5, Molecular Devices). Excitation and emission wavelengths for eGFP were set as 485 nm and 538 nm, respectively. Each measurement was conducted in Corning 96-well clear flat bottom polystyrene plates with 250 µl culture sample resuspended in 1xPBS (137 mM NaCl, 2.7 mM KCl, 4.3 mM Na_2_ HPO_4_, 1.47 mM KH_2_ PO_4_, pH 7.4). For signal normalization, raw fluorescence intensity was divided by OD _600_ values of each sample.

For 0-to-1 normalization, each value was subtracted from minimum value and divided by difference between maximum and minimum values in related groups. The data for the initial expression level was recorded as the 15 ^th^ minute to eliminate errors caused by delays in early protein expression.

### HspR Expression and Western Blot Analysis

*E. coli* BL21 (DE3) cells (New England Biolabs, Inc.) carrying HspR expression vector (T7-HspR-pET22b) was grown in LB medium with proper antibiotics at 37°C and 180 rpm in Erlenmeyer flasks. 1% of inoculums from overnight cultures were used to start expression cultures and monitored until OD _600_ reaches 0.4-to-0.6 before induction steps were applied. A control culture was kept as uninduced and other culture was induced with 1 mM isopropylthio-galactoside (IPTG) for 3 hours. Afterwards, cells were collected, resuspend in 10 mM imidazole buffer (pH 7.4) supplemented with 1 mM phenyl methyl sulfonyl fluoride (PMSF) (AMRESCO Inc.), and proteins were extracted via freeze-thaw method. Protein concentrations were determined with BCA Assay (Thermo Scientific) and diluted to final concentration of 740 µg/ml. Proteins were denatured and resolved on 15% SDS-PAGE gel prepared with BioRad SDS Gel casting system. 20 μL from protein samples were run on gel by using 6x Loading Dye (375 mM Tris-HCl (pH 6.8), 9% SDS (w/v), 50% Glycerol (v/v), 0.03% Bromophenol blue (v/v)). All samples were boiled at 95°C for 5 min prior to run on gel. 1X SDS Running Buffer (25 mM Tris-HCl, 200 mM Glycine, 0.1% (w/v) SDS) was used during run. Further, gel was transferred to polyvinylidene difluoride (PVDF) membranes (Thermo Scientific). Membrane was blocked with 5% freeze-dried nonfat milk in TBS-T for 1 h at room temperature, and then incubated at 4°C overnight with primary antibody (His-Tag mouse McAb) (Proteintech Europe) diluted at 1:10000 in blocking solution. Afterwards, membrane was washed with TBS-T and incubated with HRP conjugated goat anti-mouse secondary antibody (abcam) diluted at 1:10000 in blocking solution for 1 h and visualized by enhanced chemiluminescence (Bio-Rad) according to the manufacturer’s protocol on ChemiDoc™ Imaging System with Image Lab™ Software – Bio-Rad.

### RNA Purification and cDNA Synthesis

NucleoSpin RNA kit (New England Biolabs, Inc.) was used according to manufacturer’s instructors to isolate total RNA from each sample. Three independent biological replicas were prepared for each group. RNA concentration was quantified with NanoDrop 2000 spectrophotometer (Thermo Scientific). Samples were stored at −80°C until cDNA synthesis.

Reverse transcribed RNA concentration were set to 500 ng in 20 µl reaction volume for each sample. iScript cDNA Synthesis Kit (Bio-Rad) were used to convert RNAs into cDNAs according to manufacturer’s instructions (5 min at 25°C, 20 min at 46°C, and 1 min at 95°C).

### qPCR and Data Analysis

After cDNA preparation, qPCR experiment was performed with 1 µl cDNA for each sample via SsoAdvanced Universal SYBR Green Supermix (Bio-Rad) for *egfp* and a housekeeping gene (*hcaT*) according to manufacturer’s instructions. Primers were specified in Table S1. Three technical replicas were prepared for each independent biological replica. PCR cycles were proceeded as follows: initial denaturation for 3 min at 95°C, denaturation for 10 sec at 95°C, annealing for 15 sec at 60°C, extension for 10 sec at 72°C for 39 cycles, 10 sec at 95°C. Product specificity was confirmed by a melting curve analysis (65°C-95°C). Comparative Ct method (ΔΔCt) was used to analyze results.

### Microscopy

Samples were prepared together with each fluorescence measurement assays with specified time points in each figure. All imaging was conducted with LSM 510 Confocal Microscope (Zeiss). Samples were excited with Argon 488 nm for reporter imaging and emission was collected with LP 505 filter for eGFP while QD samples were excited with HeNe 543 nm laser and emission was collected with LP 585 filter. For dose-response curve analysis, bright field imaging of samples were also conducted which were merged with corresponding fluorescence images, afterwards.

### Statistical analysis

All data were expressed as mean ± standard error mean. Depending on the groups of interests, either one-way analysis of variance (ANOVA) or two-way ANOVA with Dunnett’s/Tukey’s/Sidak’s multiple comparison tests (GraphPad Prism v6) were used to compare groups.

## ASSOCIATED CONTENT

**Supporting Information** accompanies this paper.

A list of primers, a list of gene sequences, Sanger sequencing verification results of all vectors, plasmid maps of all vectors, Confocal microscopy images, RT-qPCR results, Western Blot result of HspR expression, growth curves of heat and QD treated samples, associated references.

## AUTHOR INFORMATION

### Author contributions

UOSS lead the study. UOSS and BS formed the experimental plan. BS, designed, and cloned the genetic circuits, ran the toxicity test and analysed the outputs of the circuits. BS and NH carried out the expression analysis of HspR protein. UOSS and BS analysed the data and wrote the manuscript.

### Notes

The authors are in the process of preparing a patent application for the HspR protein based genetic circuit.

## ACKNOWLEDGEMENTS

We thank TUBITAK for the financial support through project 114Z653. UOSS acknowledges the TUBA-GEBIP Award, BAGEP Award. We also thank Prof. Hilmi Volkan Demir for the generous gift of the Q-Dots from his laboratory. We thank Dr. Esra Yuca for technical support.

## ABBREVIATIONS

crRNA: cis-repressing RNA
HAIR: HspR-associated inverted repeats
HSP: heat shock protein
HSPR: heat shock protein response
IR2: inverted repeat 2
IR3: inverted repeat 3
QD: quantum dot
RBS: ribosome-binding site
ROS: reactive oxygen species
sncRNAs: small non-coding RNAs
taRNA: trans-activating RNA.

---

**TOC FIGURE**

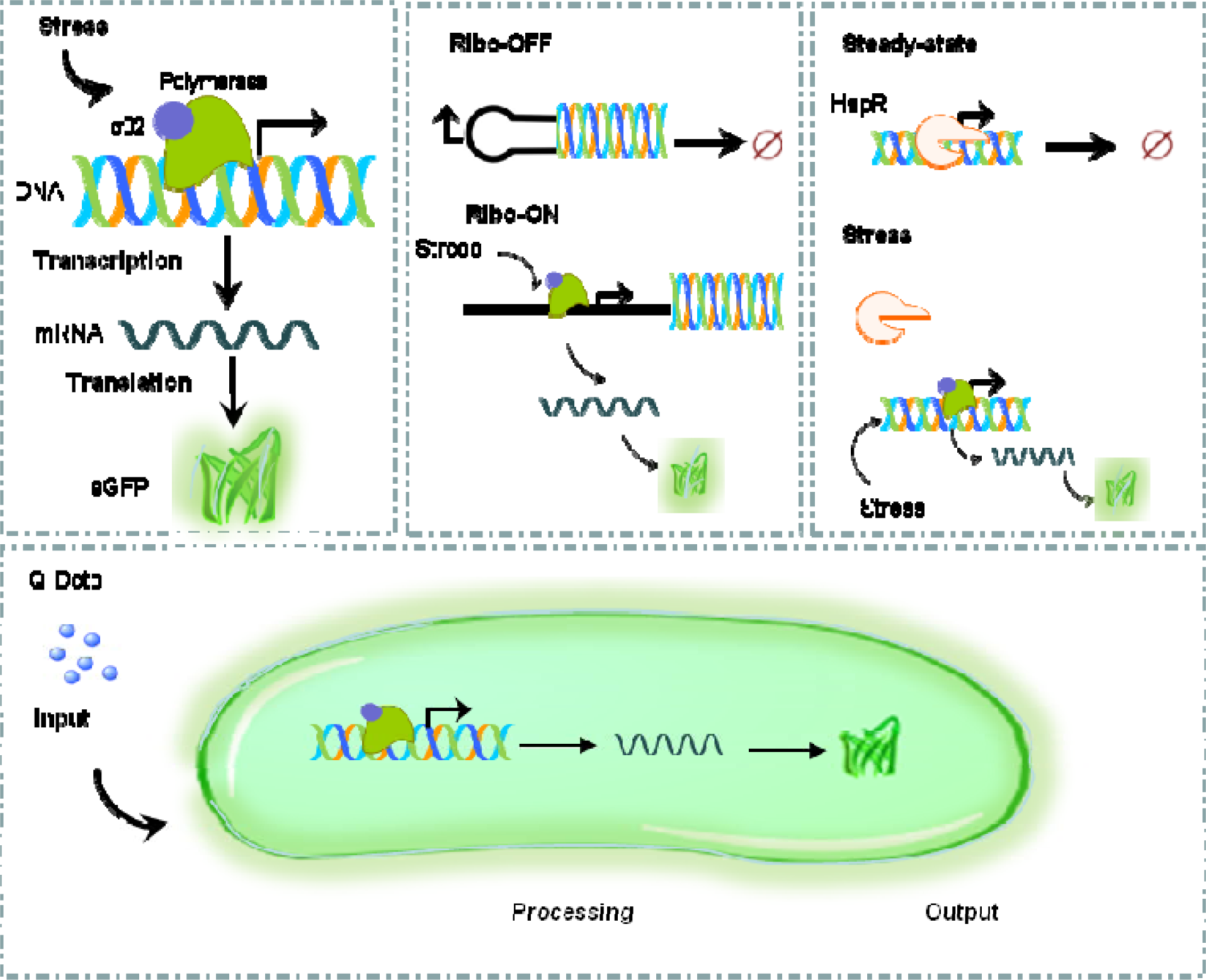

